# Probabilistic language models in cognitive neuroscience: promises and pitfalls

**DOI:** 10.1101/168161

**Authors:** Kristijan Armeni, Roel M. Willems, Stefan Frank

## Abstract

Cognitive neuroscientists of language comprehension study how neural computations relate to cognitive computations during comprehension. On the cognitive part of the equation, it is important that the computations and processing complexity are explicitly defined. Probabilistic language models can be used to give a computationally explicit account of language complexity during comprehension. Whereas such models have so far predominantly been evaluated against behavioral data, only recently have the models been used to explain neurobiological signals. Measures obtained from these models emphasize the probabilistic, information-processing view of language understanding and provide a set of tools that can be used for testing neural hypotheses about language comprehension. Here, we provide a cursory review of the theoretical foundations and example neuroimaging studies employing probabilistic language models. We highlight the advantages and potential pitfalls of this approach and indicate avenues for future research.

## 1 Introduction

Neuroimaging studies of language comprehension have over the course of the past decades generated a wealth of data which have inspired several neurobiological models (e.g., Friederici, 2012; Hagoort, 2013; Hickok and Poeppel, 2007). Such studies typically correlate or compare task-based changes in cognitive processing with changes in neural metabolic demands by means of functional magnetic resonance imaging (Logothetis, 2008) or changes in electrophysiological responses with magneto- or electroencephalography (Luck, 2005; Hansen et al., 2010). In a complementary fashion, brain stimulation techniques can be used to stimulate or perturb neural populations and thus to probe the relevant pathways for language comprehension (Devlin and Watkins, 2007).

More broadly, one of the main goals of cognitive neuroscience is to identify the explanatory relations between neuronal and cognitive computations that account for behaviour (Poldrack, 2010; Poeppel, 2012). This requires explicit formalization of the hypothesized cognitive computations (Forstmann and Wagenmakers, 2015; Palmeri et al., 2016). It has been noted before that the relative lack of well-defined computational characterizations of comprehension processes is one important factor hindering progress in explanatory understanding of the neurobiology of language (Hagoort, 2009; Embick and Poeppel, 2015).

Recently, such motivations have led to adoption of computational linguistic methods in cognitive neuroscience (Brennan, 2016). Grounded in expectation-based theories of sentence comprehension (Hale, 2001; Levy, 2008), *statistical* or *probabilistic language models* which assign conditional probabilities to linguistic representations (e.g., words, words’ parts-of-speech, or syntactic structures) in a sequence are increasingly being used, in conjunction with information-theoretic complexity measures, to estimate word-by-word comprehension difficulty in neuroscience studies of language comprehension (Figure 1).

**Figure 1:**
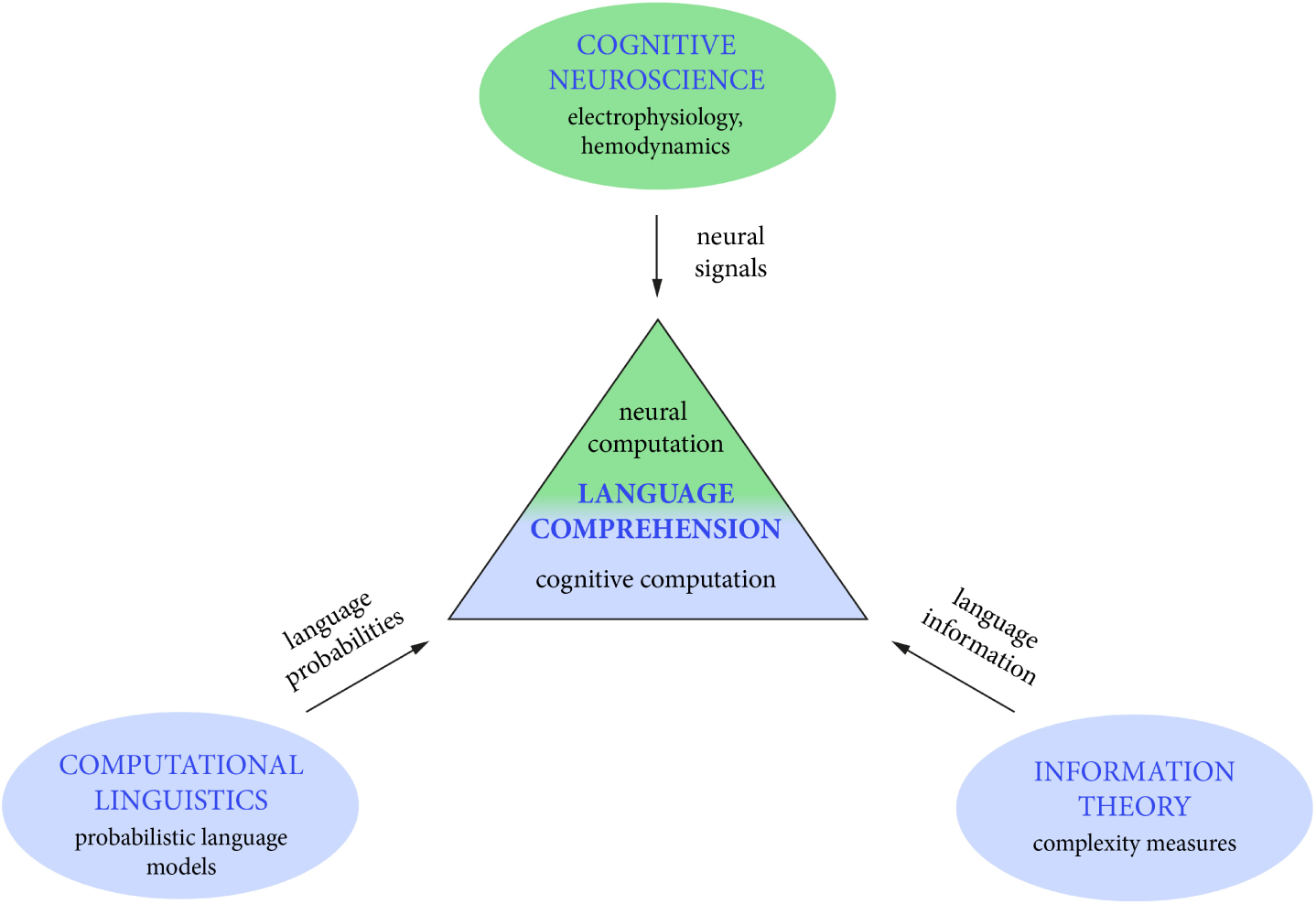
A schematic depiction of the interdisciplinary collaboration between probabilistic modeling and cognitive neuroscience of language.

While the use of probabilistic language models represents a step forward towards explicit account of expectation-based cognitive computations, it is important to critically acknowledge both the respective strengths and limitations. What are the promises and pitfalls of the approach? What can we expect to learn from it? Can probabilistic modeling go beyond localizing candidate cognitive computations in space and time?

The purpose of this paper is to provide a balanced review and discussion of the use of probabilistic language models in cognitive neuroscience of language. We first provide a cursory review of the general framework and review recent example applications. We then critically discuss the promises and limitations of this nascent interdisciplinary bridging. We conclude by outlining outstanding questions for future research.

## 2 Probabilities and information in language

### 2.1 Probabilistic constraints in language processing

Probabilistic models of cognition have witnessed a surge of interest in recent years (Chater et al., 2006). In the domain of language, it has generally been recognised that the cognitive system is sensitive to distributional properties of the language input and that probabilistic constraints play a role in both early language acquisition and later language processing (Kuhl, 2010; Griffiths, 2011; Seidenberg, 1997). Empirical support for human sensitivity to statistical/probabilistic constraints at the level of words has been shown through the word frequency effect on word recognition, disambiguation, and ease of processing (see Jurafsky, 2002, for review and evidence). Additionally, the role of statistical/probabilistic constraints in language processing and production has been shown through the effect of contextual constraints, that is, as graded sensitivity of behavioural or neural measures (e.g., reading times or amplitudes of event-related potentials) to how constraining the prior context is on possible sentence continuations (Gibson and Pearlmutter, 1998).

The effects of contextual constraints and word probabilities are commonly interpreted as reflecting some form of graded prediction, expectation or anticipation in language comprehension (Huettig, 2015; see also Kuperberg and Jaeger, 2015, for a terminological remark). Word probability in sentences is normally measured by means of human judgments in the cloze task (Taylor, 1953). In this task, participants are presented with sentence contexts where the target word position is blank. They are asked to fill the blank with a plausible word. The cloze probability of a word is then determined by counting the number of participants that used the word to continue the sentence.

Word probability effects and effects of contextual constraints provide evidence that graded statistical/probabilistic constraints in the linguistic signal and linguistic experience more broadly impact the real-time human language processing system; however, in experimental settings the exact computations explaining such effects are often not modelled explicitly. In what follows, we provide a cursory review on how probabilistic information can be modeled and quantified formally.

### 2.2 Probabilistic language models

In a nutshell, probabilistic language models are mathematical formalisms describing probability distributions over language data. One of the most common applications of probabilistic language models is in so-called sequence-prediction tasks. In the case of language, this means probabilistic models can be used for generating expectations about upcoming words given the words seen so far in a sentence (usually up to a limited length).

A distinction can be made between sequence-based language models that predict the words based on sequences of past words—a domain also called “statistical language modelling”—and models that estimate the probability of a *syntactic structure* underlying the observed sequence of words or the probability of the upcoming word given the syntactic parse so far (see also Section 2.2.1 and Figure 2 below). This is a domain proper to “computational linguistics” and as such normally considered distinct from statistical language modeling; there is, however, a great deal of overlap between the two research domains (Rosenfeld, 2000).

**Figure 2:**
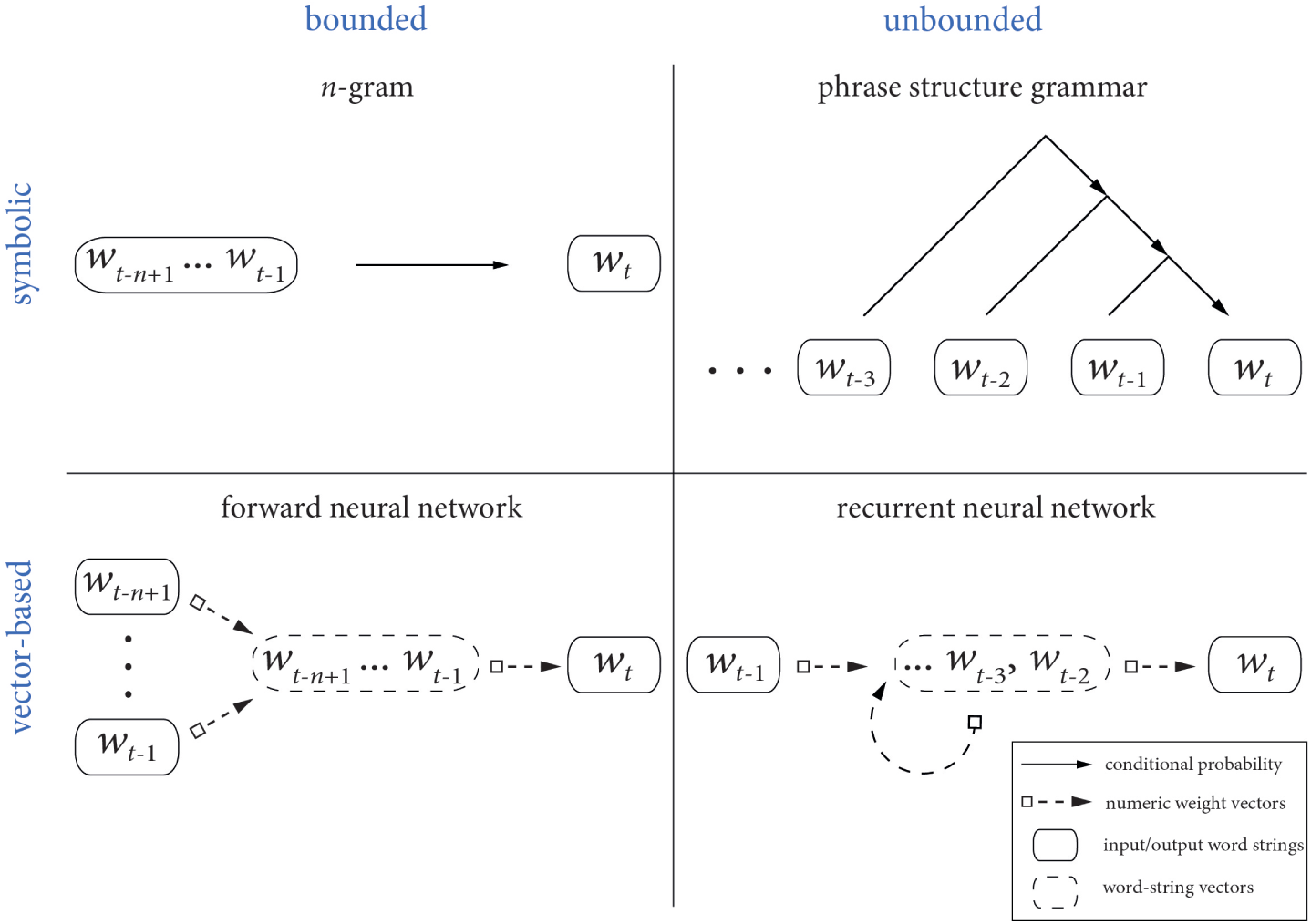
Classification of language models according to context-boundedness and the nature of representations. Classification according to the type of representation is depicted row-wise, and amount of context column-wise.

For the sake of convenience, we will in this review subsume this distinction and use a single term “probabilistic language models” because the neuroimaging studies reviewed presently in Section 3 employ descriptions at both levels of granularity.

#### 2.2.1 Common ways of estimating probabilities

How are language probabilities estimated? Three broader classes of models are commonly used in computational psycholinguistics: *n*-gram models, phrase structure grammars (PSGs), and neural networks.

*N*-gram models, also known as Markov models, represent the simplest architecture for estimating the probabilities. The term *n*-gram stands for any sequence with the length of *n*-items where the model order *n* denotes the number of context words (*n* – 1) plus the word (*n*-th word) for which the probability is computed (Jurafsky and Martin, 2009). Therefore, a 4-gram model takes into account three preceding words in a sequence for computing the conditional probability of occurrence for the fourth word. The basis for computing these probabilities are the relative frequencies of co-occurrence of word sequences derived from the training data in language corpora. We add that an *n*-gram can stand for the sequence of actual words or, alternatively, syntactic categories of words (or *parts*-*of*-*speech*).

Apart from *n*-grams, probabilities of the upcoming words can also be estimated by using either feed-forward (FNN) or recurrent neural network (RNN) architectures (Bengio et al., 2003; see, De Mulder et al., 2015, for a recent review on RNNs). In these architectures, the words are not represented as symbol strings as with *n*-grams, but are instead converted into *vector* representations; each word is coded as a sequence of real numbers—a real-valued feature vector. These vector representations are given as input to a pre-specified number of neuron-like hidden units where activation of these units is given by mathematical functions and transformations applied to the word vectors.

In recurrent neural networks, the hidden units also receive recurrent input from the states encoded in previous steps (see Figure 2, bottom right) which means any current state of the layer reflects the history (of an undetermined length) of past network states (e.g., representing sentential context in language tasks). During model training in word prediction tasks, the models adjust the weights (or parameters) assigned to each hidden unit and individual components of word vectors such that the difference between words predicted by the model and words that actually appear is minimized. The activation of the output units are rescaled such that the output vector can be interpreted as a probability distribution over words. Each unit’s activation is the estimated probability that the corresponding word will appear next, given the word sequence presented to the model.

PSGs are sets of so-called *rewrite rules* relating a phrasal class (e.g., a noun phrase) to its constituent parts of speech (e.g., a determiner, an adjective, and a noun) to the actual word strings (e.g., “a red hat”). A PSG therefore provides, by sequentially applying the rewrite rules in a process called *derivation*, the structural description underlying a given sequence of words. A *probabilistic* (or *stochastic*) PSG assigns a probability of a syntactic parse given a surface level string, or the probability of the upcoming word given the syntactic parse so far (Roark 2001, see also Figure 2, top right). The probability of the entire parse is determined as a joint probability of all rewrite rules needed to generate the complete parse. The probabilities of rewrite rules are determined from occurrences in syntactically-annotated corpora known as *tree*-*banks* (see, e.g., Marcus et al., 1993).

#### 2.2.2 Context-boundedness and representations

To see how these classes of models compare to one another, it is useful to consider their characteristics along two key dimensions: whether there is a limit to the amount of context considered for computing the conditional probabilities (boundedness) and what is the nature of representations over which these models compute. This gross classification with boundedness and representations is schematically depicted in Figure 2.

In terms of the amount of context that can be taken into account for estimating the probabilities, models fall either in the category of *bounded* or *unbounded* models (represented column-wise in Figure 2). Bounded models impose a finite bound to the length of the preceding context considered; model classes with bounded limit are the *n*-gram models and feed-forward neural networks where the probabilities are conditioned on a fixed number of preceding words. Recurrent neural networks and PSGs, on the other hand, are unbounded models. A recurrent neural network’s hidden layer activation depends on the entire input string so far (Figure 2, bottom right) whereas in PSGs the current word can depend on words at *any* earlier point which makes it possible to model long-distance dependencies between the words—a hallmark of language structure.

The second classification dimension is the nature of representations over which the models compute (represented row-wise in Figure 2); specifically, the representations can either be *symbolic* or *vector*-based (the latter are also termed as: analogue, continuous or distributed representations). *N*-grams and PSGs fall into the first category whereas feed-forward and recurrent neural networks operate over continuous or distributed vector word representations. The critical difference between the two types of representations is that symbolic representations (e.g., word strings “dog” and “cat”) can only be equal or unequal with no inherent measure of similarity apart from the relationship reflected in the frequency of co-occurrence; in contrast, numerical, vector representations in neural networks can be compared using a similarity measure. For example, because a every vector has a direction in a vector-space, a distance between two word vectors (quantifying semantic distance between two words encoded by these vectors) can be computed mathematically as a function of the angle between two vectors (smaller angle indicates more closely related words).

### 2.3 Quantifying complexity: entropy and surprisal

On the basis of probabilities estimated with probabilistic models described above it is possible to compute the amount of information conveyed by each word in a sequence. This is quantified with information-theoretic complexity metrics such as word *surprisal* and word *entropy.* A complexity metric is any measure quantifying hypothesized processing difficulty at the current word and need not be probabilistic; the number of nodes traversed in a hierarchical syntactic derivation is another example of a metric capturing comprehension difficulty (Gibson and Thomas, 1999). For a complete treatment on information-theoretic complexity metrics specifically, we point the reader to a recent review by Hale (2016); here, we provide a brief overview to establish the necessary coherence with rest of the paper.

Surprisal is an information-theoretic measure quantifying how unexpected and thus how informative the current word (*w_t_*) is given the words that precede it (*w*_1_, …, *w*_*t*–1_). A higher word surprisal values indicates that the currently encountered word is less expected given the context. In mathematical terms, surprisal *S*(*w_t_*) is defined as the negative logarithm of the word’s conditional probability of occurrence:

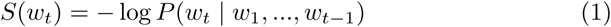

If base-2 logarithm is used, surprisal is expressed in *bits.* The same is true for the word entropy information measure, which quantifies how narrow or spread-out the probability distribution of possible next words is. If taken as a measure of cognitive effort, it models the degree of the listener’s or reader’s uncertainty about the upcoming word given the words encountered so far. Higher entropy values represent a higher degree of uncertainty (due to a higher number of possible candidate continuations) whereas lower entropy values signify a higher degree of certainty with fewer, highly probable continuations given the context so far. Mathematically, entropy at the current word position *H*(*t*) is defined as the expected value of surprisal for the upcoming word (*w*_*t*+1_) given the words encountered so far (*w*_1_, …, *w_t_*):

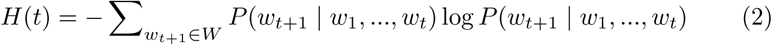

where *W* denotes the set of all possible words.

Above, we introduced surprisal and entropy as defined over actually observed words in sentences, however, both metrics can also be computed on the basis of words’ parts-of-speech (Frank, 2010) or syntactic structures as obtained from probabilistic grammars (Hale, 2003; Roark et al., 2009). If the models take into account the actually observed words, a metric is said to be *lexicalized*, whereas in the case of *unlexicalized* metric, only structural probabilities or probabilities of parts-of-speech are used for computing complexity (Demberg and Keller, 2008). In other words, unlexicalized complexity metrics are not concerned with lexical-semantic properties of language input. However, additional assumptions are required on the type of syntactic structures plausibly involved in human comprehension (Hale, 2003; Frank, 2013).

In addition to surprisal and entropy, another relevant complexity metric is *entropy reduction.* Originally, Hale (2006) defined the entropy reduction resulting from integrating word *w_t_* into the derivation of the sentence so far, as the amount by which uncertainty about the complete sentence’s structure gets reduced by excluding structures incompatible with *w_t_*. In practice, however, estimating the probabilities of all possible sentence structures is not feasible. For this reason, the scope of the entropy computation has been reduced to, for example, the possible sentence continuations (Wu et al., 2010), a subset of upcoming four words (Frank, 2013), or even just the single next word (Roark et al., 2009).

In brief, cognitive neuroscience and probabilistic language modeling conceptually share a common point in emphasizing information processing and probabilistic aspects of language comprehension. We now turn to the literature where probabilistic language models were used to analyze neural measures of interest.

## 3 Example applications

Until recently, probabilistic language models were predominantly tested against behavioral data, such as grammaticallity judgments, self-paced reading times, and eye-movements (e.g., Boston et al., 2008; Demberg and Keller, 2008; Frank and Bod, 2011; Linzen and Jaeger, 2014; Lau et al., 2017). The use of probabilistic language models in cognitive neuroscience of language comprehenension represents a recent trend; here we review six example studies where probabilistic language models were used word-by-word to quantify complexity in sentence or story comprehension tasks. We begin by reviewing studies where information measures represented the predictor of interest and continue with those where they were used as an additional predictor to non-probabilistic complexity measures.

### 3.1 Information measures as the predictor of interest

Given that word surprisal and entropy quantify different aspects of the incoming linguistic signal, Willems et al. (2016) used 3-gram language models and asked whether the two measures yield distinct loci of activation in the brain while participants listened to auditory narratives. Word entropy negatively correlated with blood oxygen level dependent (BOLD) signal in the right inferior frontal gyrus, the left ventral premotor cortex, left middle frontal gyrus, supplementary motor area, and the left inferior parietal lobule whereas word surprisal showed positive correlations bilaterally in the superior temporal lobes and in a set of (sub)cortical regions in the right hemisphere (see Figure 3 below). These results were interpreted within the predictive coding framework; regions sensitive to entropy were taken to reflect active predictions of the coming words (predictions are possible in low entropy states) and areas related with word surprisal (how surprising the current word is) were interpreted as possibly reflecting prediction errors in the early auditory areas.

**Figure 3:**
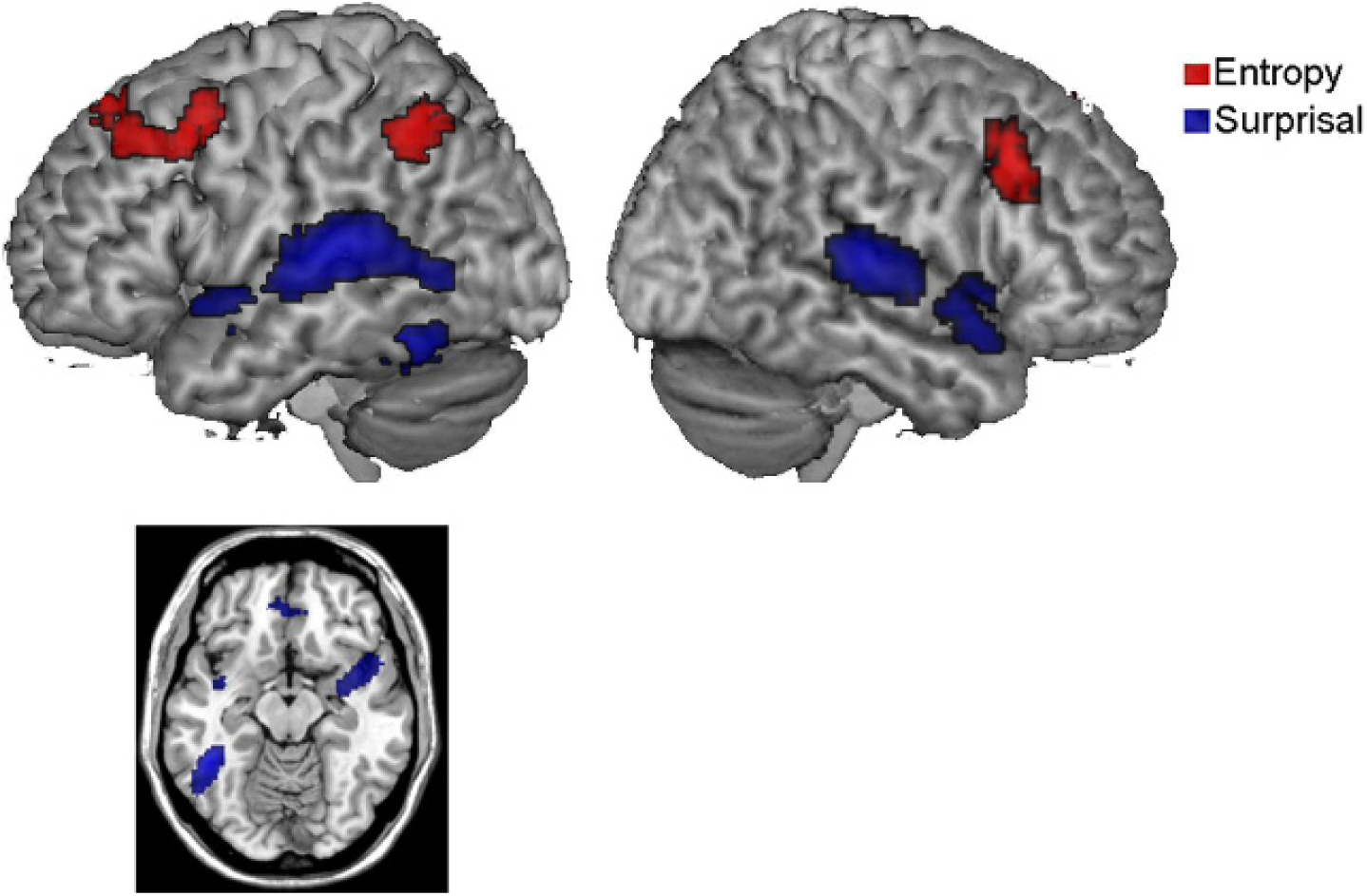
Brain areas activated more strongly (real stories compared with reversed story fragments) for word surprisal (blue) and word entropy (red). Reproduced from Willems et al. (2016).

As explained in sections 2.2.1 and 2.3, language probabilities and complexity metrics can also be computed on the basis of syntactic structures. Henderson et al. (2016) used the probabilistic phrase structure parser by Roark (2001) to study the cortical infrastructure sensitive to syntactic surprisal during naturalistic comprehension. The authors simultaneously measured BOLD responses and eye-movements while participants silently read stories in paragraphs. A whole-brain comparison between word groups with high and low syntactic surprisal revealed significant differences in the inferior frontal gyrus bilaterally, left anterior temporal lobes (under a less conservative statistical threshold), bilateral insula, fusiform gyrus, and the putamen. There were no statistically significant predicted differences in superior temporal lobes or the superior temporal sulcus.

The authors discuss the results as in line with current neurobiological models that place the cortical systems for syntactic computations to inferior frontal and anterior temporal cortices. It is interesting to note that eye-tracking data revealed no differences for the syntactic surprisal contrast; this stands in contrast to previous reports showing relations between syntactic surprisal metrics and eye movements (e.g., Boston et al., 2008; Demberg and Keller, 2008). The authors speculate that the novel use of a lexicalized syntactic surprisal—as opposed to unlexicalized syntactic surprisal used in previous reports—might be a possible source of discrepancy.

In cognitive electrophysiology, one of the most studied signals is the event-related potential (ERP); time-averaged voltage deflections reflecting an integrated (summed) response of large populations of spatially and temporally coherent cortical pyramidal neurons (Luck, 2005). Under the assumption that those models and complexity metrics that best explain the data also more closely resemble putative cognitive mechanisms, Frank et al. (2015) computed word surprisal and entropy reduction of words and their parts-of-speech under three types of models: *n*-grams (*n* = 2, 3, and 4), phrase-structure grammars, and recurrent neural networks.

Out of all the possible relations between word information measures and six candidate ERP component amplitudes from an exploratory analysis, word surprisal measure computed on the basis of 4-grams and RNNs significantly improved the fit of the regression model to the N400 ERP amplitude over and above PSGs but not vice versa; that is, the inclusion of hierarchical syntactic information in the models was not reflected in better statistical fit. In terms of mechanistic interpretation, the authors take this result as compatible with the lexical retrieval account of the N400 component (Kutas and Federmeier, 2000).

### 3.2 Information measures as additional predictor

The studies reviewed above looked exclusively at the effects of information measures computed by probabilistic language models. We now turn to studies where such measures are investigated in addition to non-probabilistic measures of complexity.

Brennan et al. (2016) investigated the neural correlates of syntactic complexity during naturalistic comprehension. Comprehension difficulty was characterized with *n*-grams, PSGs, and minimalist grammars (a formal grammar that accounts for syntactic phenomena not accounted for by PSGs). A stepwise inclusion of progressively more “syntactically sophisticated” language predictors improved the statistical fit to BOLD time courses in the bilateral anterior temporal lobes, left inferior frontal gyrus, left posterior temporal lobe, left inferior parietal lobule, and left premotor area. When taken on their own, the 2- and 3-gram surprisal measures revealed significant effects in the anterior temporal lobes, left inferior frontal gyrus and the left posterior temporal lobe.

Based on the fact that models including knowledge of hierarchical syntax explained variance over and above the models that incorporate only linear, word sequence-based statistics, the authors take their results as evidence for the involvement of abstract syntactic linguistic knowledge in every-day sentence comprehension. The effects of surprisal are in part consistent with the results by Willems et al. (2016) who similarly report word surprisal effect in the posterior temporal lobe.

Nelson et al. (2017) investigated modulations of average high frequency (70150 Hz) power in intracranially recorded electrophysiological signals by hypothesized syntactic phrase-structure building operations during a word-by-word sentence reading task. In model-comparison analysis, they contrasted explanatory power of non-probabilistic hierarchical syntactic predictors (counting the number of open syntactic nodes at the moment when each word was presented) and probabilistic language models. The former showed significant effects in several superior temporal and inferior frontal electrode sites, whereas lexical and part-of-speech bigram surprisal (i.e., transition probability) and next-word entropy showed positive and negative effects, respectively, in electrodes surrounding the middle temporal gyrus.

Based on these results, the authors argue in favor of neurophysiological reality of hierarchical syntactic operations during comprehension. They interpreted the probabilistic predictability effects as consistent with other reports localizing neural generators of single-word semantic priming, N400, and repetition suppression effects to posterior temporal regions.

Van Schijndel and Schuler (2015) investigated the role of syntactic memory load during auditory story comprehension. The strength of spectral coherence of MEG oscillatory neural activity in the 10 Hz range was taken as a neural indicator of increased working memory usage. Syntactic complexity was quantified as the number of incomplete syntactic structures maintained at any word position (depth of syntactic embeddedness estimated based on the most likely parse of a probabilistic PSG). *N*-gram probability predictors and a PSG surprisal were used as control measures.

The authors report that the average alpha-band coherence in a pair of left posterior and anterior sensors range was significantly different for two levels of syntactic depth while controlling for *n*-gram probability effects; trigram probability showed marginal alpha coherence effects prior to correcting for multiple comparisons. Similar to the interpretations by Brennan et al. (2016) and Nelson et al. (2017), the authors interpreted the results as showing that hierarchical linguistic structure is computed during comprehension because it improves the fit to empirical data over competitive non-hierarchical models.

Finally, apart from regression-based analyses and factorial designs, the statistical relationship between neural data and language model output can also be ascertained by means of multivariate statistical techniques, for example, by using features of a language model in an intermediate step for decoding stimulus identity from multivariate neural data. Wehbe et al. (2015) report that binary word classification accuracy based on MEG amplitudes, which in turn were predicted by RNN output vectors—interpreted as word probabilities—, was highest approximately 400 msec after word onset, which can be seen as consistent with results by Frank et al. (2015) who found a positive correlation between lexical surprisal and the N400 amplitude. On the basis of the time-course of classification accuracy, the authors linked the late effect of word probability to word integration processes (that differ between unpredictable and predictable words).

### 3.3 Summary

Current applications of probabilistic language models in cognitive neuroscience show that probabilistic language models can be used with hemodynamic and electrophysiological methods and allow researchers to investigate and focus on spatial fingerprints for specific linguistic computations in cortical regions (Willems et al., 2016; Henderson et al., 2016) or to compare predictions of different models against each other on the basis of same neurobiological data, be it fMRI time courses (Brennan et al., 2016), language event-related M/EEG components (Frank et al., 2015; Wehbe et al., 2015), or spectral contents of electrophysiological signals (Van Schijndel and Schuler, 2015; Nelson et al., 2017). The studies employed language stimuli in both auditory and visual modalities and, with the exception of the studies by Frank et al. (2015) and Nelson et al. (2017), used language stimuli in naturalistic, narrative contexts. We now turn to a more detailed discussion of specific advantages and disadvantages of the approach.

## 4 Advantages

### 4.1 Formalized cognitive computations

What can we expect to learn from model-based analyses? Probabilistic language models represent the computational level of explanation in cognitive neuroscience in the time-honoured sense of Marr (1982): What aspect of the language input enters into the computation? What is being computed and why? Quantitative methods represent a complement to subtraction paradigms in neuroimaging (see Hagoort, 2014, for a recent review on sentence comprehension) where cognitive computations are inferred on the basis of informal, qualitative task-based cognitive contrasts.

Reading off cognitive computations from tasks is not straightforward (Boone and Piccinini, 2016) in that it must first be assured that the task taps into the target linguistic computation and not, for example, meta-linguistic processes. This can be assured by comparing several informal task contrasts (see, e.g., Kaan and Swaab, 2002, for a discussion on task contrasts for syntactic computations) or by computationally modelling the task itself (see, e.g., Norris et al., 2000, for a model of phoneme monitoring). Only once this is established, it is possible to draw links to the observed neural effects. In model-based approaches, however, markers of sentence-level cognitive computations, for example syntactic surprisal, are directly statistically related to neural signals.

From a methodological perspective, explicit mathematical definitions and computational implementations lead to a more rigorous and standardized quantification of independent variables which reduces dependence on researchers’ operationalizations of specific concepts (but see section 5.1 for potential pitfalls related to allures of formalization).

### 4.2 Theory evaluation

In other domains of cognitive neuroscience, such as decision-making and cognitive control, linking neural data to parameters of formal models served as a fruitful way to overcome the impasse when competing models could not be distinguished based on overt behavioral responses alone (Forstmann et al., 2011). As such, model comparison proved to be a major contribution of combining model-based approaches with neurobiological data (Mars et al., 2012). Given the fast pace and incrementality of language comprehension processes, *covert*, online measures of comprehension difficulty such as eye-movement records have been a key component of empirical evaluations for competing models in reading and spoken comprehension (see Rayner, 1998; Huettig et al., 2011, for reviews).

Brain signals can be similarly considered as covert markers of online cognitive difficulty and as such taken as empirical test bed for cognitive hypotheses implemented in language models. Whereas compared to model-based approaches in other domains, neural measures do not necessarily represent exclusive diagnostic data for evaluating cognitive theories, any neurophysiologically valid cognitive theory should ultimately account for neural measures as these are closely linked to the underlying neural computations. As such, neural validation of cognitive theories provides cognitive-computational constraints for plausible neuronal computations (Mars et al., 2012; Palmeri et al., 2016, see also Section 5.5 below).

### 4.3 Statistical efficiency in analyses

In most current empirical applications of language models, complexity metrics are computed for all words in experiments which improves statistical sensitivity in the studies compared to the traditional experimental approach. For example, the three stories used by Willems et al. (2016) yielded approximately 3,000 words, all of which were considered as separate trials in the analysis. This contrasts with the currently prevailing experimental approaches, where, most often, studies will only investigate neurobiological effects on target words in non-filler items. This of course follows from the logic of experimental designs; however, it also means that large stretches of neural data are collected without being inspected or considered in the analysis.

Further, probabilistic language models provide a quantification over a range of values, rather than only the extreme poles of the spectrum which is common in subtraction-based designs (but see, e.g., Pallier et al., 2011, for an exception). In case of significant statistical dependence between variables, parametric variation gives stronger support to the actual workings posited by the model compared to factorial designs (Bechtel and Abrahamsen, 2010).

### 4.4 Naturalistic stimuli and data reuse

Apart from explicitness and increased statistical sensitivity of research designs, there is another potential advantage of language modelling: it makes it easier to study the brain responses to naturalistic stimuli (Brennan, 2016). Even though the study of language in its ecological setting has in certain cognitive traditions been regarded as an ill-advised enterprise on principled and practical grounds (Chomsky, 1959, 1995), it was highlighted as a necessary empirical step to study the brain from the systems level (see Hasson and Honey, 2012; Small and Nusbaum, 2004).

The approaches reviewed here strike a balance between the two perspectives: while the computational part enables rigorous formalization of the cognitive hypothesis, absence of secondary task during the experiment enables the study the of brain responses to more ecologically valid stimuli. Studying the brain in naturalistic settings is a desirable research approach (for a recent overview of challenges and developments, see the contributions in Willems, 2015), nevertheless, we hasten to add that it should complement established experimental approaches which capitalize on well-controlled task-based designs (see e.g. Fetsch, 2016, for a recent opinion on the importance of experimental designs); for example because a specific cognitive hypothesis might not be available and implemented as a probabilistic language model.

It is also worth emphasizing that the absence of specific task constraints in the experimental design lends these types of neuroimaging data sets appropriate for reuse and sharing for analyses with new language models that embody novel hypotheses; a component of contemporary research practice which is being actively recognized in the neuroscience community (Poldrack and Gorgolewski, 2014).

## 5 Limitations and pitfalls

Even though probabilistic modeling comes with evident advantages, it has, as is true for any methodological advancements, specific limitations. In light of increasing acceptance of model-based analyses by experimental cognitive neu-roscientists, it is important to render these pitfalls explicit.

### 5.1 Allures of formalization

Due to their computational implementation and quantitative nature, formally estimated language probabilities can be seen as representing a more objective estimate than measures of cloze probability obtained on the basis of subjective, human judgments (Staub, 2015). It is true that language models and complexity metrics improve the comparability between experiments and can be viewed as more objective from that point of view.

Nevertheless, even for formal estimates the extent to which they capture the “ground truth” can be debated. Using complexity measures obtained from a single language model on experimental stimuli would be comparable to using judgments of a single participant for quantifying measures of cloze probabilities (see also Smith and Levy, 2011, for discussion on the two types of language probabilities). The complementarity of the two ways of estimating probabilities is further underscored if we consider that in speech recognition tasks, for example, human judgments (providing knowledge not captured in the models alone) can be used to *improve* model performance (Rosenfeld, 2000).

Second, probabilistic language models describe the probability distributions over words but do not model the human language acquisition trajectory. Specifically, models are trained on large amounts of language data which does not correspond to how such knowledge is acquired by humans, who exploit a variety of other multimodal sensory and social cues (see Kuhl, 2010; Saffran, 2003, for reviews). From an explanatory perspective, it would therefore be inaccurate to implicitly treat models trained on collections of text as models of language acquisition.

### 5.2 Lexical confounds

All ways of estimating formal language probabilities, in one way or another, rely on observed *frequencies of occurrence* in collections of texts – language corpora. Together with the fact that complexity measures are computed on a word-by-word basis, this means that by construction probabilistic complexity measures are likely to correlate with well-known lexical nuisance variables in psycholinguistics, for example, lexical frequency (i.e., unigram probability), word length, phonological neighbourhood size, transitional probability (i.e., bigram probability, etc.).

These lexical measures characterize separate aspects of words. For example, lexical frequency is a property of the word alone whereas a 4-gram probability is conditioned on the three preceding words and therefore operationalizes context-dependent computations. Whereas both can be viewed as effects of “lexical predictability”, they can be related to distinct cognitive computations; for example, genuine predictive processing or ease of lexical retrieval (Staub 2015; Huettig 2015, see also Kuperberg and Jaeger 2015).

Given that probabilistic language models afford the use of less experimentally constrained, naturalistic stimuli, confound variables must be controlled statistically. They should be included as covariates of no interest in regression-type analyses; for example Frank et al. (2015) included word frequency, word length, and word position in the sentence as nuisance variables. Alternatively, in factorial designs, it must be ensured that experimental conditions are chosen such that they are matched for other lexical variables as was done in Henderson et al. (2016). The list of potentially confounding variables can extend depending on the experimental settings; in an eye-tracking study, Demberg and Keller (2008), for example, included also the eye-movement specific variables about whether the previous word was fixated or not, launch distance, and fixation landing position in addition to word length, word frequency, forward transitional probability, backward transitional probability, and word position in the sentence.

### 5.3 Syntactic and semantic complexity

A distinction between abstract, syntactic computations and meaning-bearing semantic operations has been a cornerstone in cognitive sciences of language and represents a theoretical framework for research cognitive neuroscience (see, e.g., Friederici and Weissenborn, 2007; Kuperberg, 2007, for discussion). A word’s frequency of co-occurrence is in principle governed by both its syntactic valence and its lexical-semantic relationships to neighbouring words. In terms of probabilistic language models, it is important to note that lexical, word-based probabilistic language models (*n*-grams, RNNs) reviewed presently cannot disentangle sources of semantic and syntactic complexities apart.

Whereas the issue of resolving semantic and syntactic influences at the level of words seems to be a technical rather than a principled one (see Padó et al., 2009; Frank and Vigliocco, 2011, for suggestions on formalizing syntactic versus semantic probabilities), at present, lexical-semantic influences on probability estimates can be overcome by using predictors based on unlexicalized complexity measures on the basis parts-of-speech *n*-gram models as in Frank et al. (2015) or probabilistic PSGs rather than actual words themselves as was done by Henderson et al. (2016); Brennan et al. (2016)

### 5.4 Linguistic levels of analysis

A hallmark of linguistic analyses is to view the language system as comprising of different levels of linguistic granularity, minimally of the phonological, lexical-semantic (word-based) and syntactic linguistic levels (Jackendoff, 2002). One of the important properties of language models and complexity metrics is that in practice these can be computed per each word in a sentence capturing the incrementality of human sentence processing (Hale, 2016).

However, it must be emphasized, that neural effects are cannot always be assessed for all individual words. For example, temporal evolution of the BOLD-response as measured with fMRI is slower than the presentation rate of words. However, this limitation can be overcome for instance by performing linear regression with a regressor which differs on a word-by-word basis such as perplexity or lexical frequency, (see Yarkoni et al., 2008, for illustration of this approach).

### 5.5 Explanatory status: maps or mapping?

Finally, it is worth touching upon the *explanatory scope* of the approach presented here. What constitutes an adequate account of explanation (in the sense of Craver, 2007) in cognitive neuroscience and how to approach it remains a debated topic and has received increased attention in cognitive neuroscience communities recently (see Pulvermüller et al., 2014; Embick and Poeppel, 2015; Jonas and Kording, 2017; Krakauer et al., 2017, for some recent discussions). It has been emphasized previously that localizing specific cognitive computations to circumscribed cortical areas does not in itself constitute a sufficient explanation (Poeppel, 2012).

Seeking a fit between probabilistically modeled cognitive states and neural data by means of a statistical model remains silent on the algorithmic and the neural levels of explanation. Specifically, complexity metrics are estimators of comprehension difficulty and can provide evidence for or against cognitive theories to the extent that the latter provide distinct predictions on where in a sentence the human cognitive system will experience difficulties (Martin, 2016). Currently, probabilistic models do not offer explanations in terms of *how* the cognitive (and neural) computation is achieved (but see Hale, 2011, for an algorithmic proposal). Clearly, any empirical success of probabilistic language models in explaining neural signals does not entail that mathematical formalisms, information measures or language probabilities *per se* are instantiated in the brain (Jurafsky, 2002).

From the perspective of neurophysiological explanation, current fMRI-based applications stay within what has been dubbed the “cartographic imperative” (Poeppel, 2012) with the goal of tentatively localizing hypothesized computations to gross-level brain areas (as in Willems et al., 2016; Henderson et al., 2016). On the other hand, electrophysiological results are predominantly informing cognitive theories (as in Frank et al., 2015; Van Schijndel and Schuler, 2015). However, it is becoming increasingly clear in cognitive and systems neurosciences that brain signals are not only indices representing diagnostic evidence for theories cast at the cognitive-computational levels of analyses, but are *biophysically meaningful* signals reflecting underlying neuronal computations and circuit configurations (Cohen, 2017) occurring at lower levels of spatio-temporal cortical organizations (this is conveyed by the upper part of our schematic in Figure 1). In this respect, electrophysiological methods represent a powerful tool, compared to hemodynamic methods, due to a closer link between electrophysiological events at lower spatial scales (as in Nelson et al., 2017, where high frequency power is taken to reflect neural computation).

Although model-based analyses reviewed above can reveal what information content during comprehension makes a difference in terms of neural signals, this type of correlational “bridging” represents an initial step towards a more ambitious goal of describing the plausible neural computational principles that explain the *mapping* to hypothesized linguistic/cognitive computations and taxonomies (Dehaene et al., 2015; see also Marcus et al., 2014). If probabilistic computations at some level represent a valid cognitive hypothesis underlying the behaviour, this should provide constraints on the target neural computations, mechanisms and algorithmic descriptions. Before concluding, we outline below some outstanding challenges that deserve further attention in the future.

## 6 Future challenges

Cognitive neuroscience shows that human listeners can integrate several sources of information to interpret an utterance (Hagoort and van Berkum, 2007). This translates into a long-standing challenge in the language modeling community: how can we bring probabilistic models to bear on larger linguistic units and contextually relevant information, for example by making use of discourse coherence in models of sentence comprehension, long short term memory neural networks etc. (e.g., Dubey et al., 2013; Hochreiter and Schmidhuber, 1997)?

Similarly, different classes of models perform with different success rates on empirical data. If a certain class of models (e.g., *n*-grams or PSGs) turns out to be consistently more successful empirically, what are the consequences for neurocognitive theories? Which aspect of the model architecture (the underlying cognitive hypothesis) or model training yields this difference compared to other models?

Theoretical and empirical investigations in psycholinguistics and cognitive neuroscience show that language processing consists of distinct representational and temporal scales, including, but not limited to, at the level of phonemes, words, sentences, and discourse (Jackendoff, 2002; Lerner et al., 2011). Typically, these stages are investigated in separate experiments with different experimental paradigms. Can probabilistic language models be used as a tool for investigating expectation-based processing at distinct representational and temporal levels of complexity concurrently in a single experiment within the same dataset (e.g., Lopopolo et al., 2017)?

Regardless of the specific computational theory embedded in the models, efforts should be spent in laying out the constraints to algorithmic and neurophysiological explanations (see Embick and Poeppel, 2015; Martin, 2016). How does probabilistic cognitive computation relate to the general principles of cortical organization for language and other cognitive-perceptual systems (e.g., Battaglia et al., 2012; Friederici and Singer, 2015)? What general property of cortical circuitry is required to explain any observed correlations and directions of the effects between probabilistic computation and neurobiological signals? More specifically, what neuronal circuit configuration and computation allows us to make a linking hypothesis to probabilistic cognitive computation? What statistical learning mechanisms must be in place to account for development of probabilistic computation in language (as in Kumaran et al., 2016)?

Lastly, probabilistic language models reduce the dimensions of language comprehension by focusing on the properties of the linguistic signal alone. An important explanatory consideration of the *what* and the *why* of probabilistic language computation will eventually have to account for the pragmatic and communicative perspective on language understanding: What purpose would probabilistic language computation serve in models of pragmatic language understanding as probabilistic inference (Goodman and Frank, 2016)? What does probabilistic computation entail for the rapid and flexible human communicative behaviour in social and interactional settings (see e.g., Levinson, 2015; Stolk et al., 2015)?

## 7 Conclusion

In the present paper, we provided a general overview of probabilistic language models, presented example applications in neuroscience studies, and discussed advantages and disadvantages. The approach advocated here should be viewed as complementary to the established experimental paradigms in cognitive neuroscience. Probabilistic language models provide computationally implemented tools for evaluating cognitive theories on neural data, mapping cognitive computations to gross-level brain areas, and offer tentative cognitive-computational explanation of electrophysiological responses. Future challenges lie in widening the scope of language models to meet the known characteristics of human linguistic-communicative capacities and moving from brain mapping to linking specific cognitive explanations of macroscopic brain signals to plausible underlying neuronal computations.

## Acknowledgments

The work presented here was partly funded by the Netherlands Organisation for Scientific Research (NWO) Gravitation Grant 024.001.006 to the Language in Interaction Consortium and Vidi Grant 276-89-007 to Roel Willems.

## Conflict of interests

none declared.

## References

Battaglia, F. P., Borensztajn, G., and Bod, R. (2012). Structured cognition and neural systems: From rats to language. Neuroscience and Biobehavioral Reviews, 36(7):1626–1639.

Bechtel, W. and Abrahamsen, A. (2010). Dynamic mechanistic explanation: Computational modeling of circadian rhythms as an exemplar for cognitive science. Studies in History and Philosophy of Science Part A, 41(3):321–333.

Bengio, Y., Ducharme, R., Vincent, P., and Janvin, C. (2003). A neural probabilistic language model. The Journal of Machine Learning Research, 3:1137–1155.

Boone, W. and Piccinini, G. (2016). The cognitive neuroscience revolution. Synthese, 193(5):1509–1534.

Boston, M. F., Hale, J. T., Kliegl, R., Patil, U., and Vasishth, S. (2008). Parsing costs as predictors of reading difficulty: An evaluation using the Potsdam Sentence Corpus. Journal of Eye Movement Research, 2(1):1–12.

Brennan, J. (2016). Naturalistic sentence comprehension in the brain. Language and Linguistics Compass, 10(7):299–313.

Brennan, J. R., Stabler, E. P., Van Wagenen, S. E., Luh, W.-M., and Hale, J. T. (2016). Linguistic structure correlates with temporal activity during naturalistic comprehension. Brain and Language, 157-158(June/July):81–94.

Chater, N., Tenenbaum, J. B., and Yuille, A. (2006). Probabilistic models of cognition: conceptual foundations. Trends in Cognitive Sciences, 10(7):287–91.

Chomsky, N. (1959). A review of B. F. Skinner’s Verbal Behavior. Language, 35(1):26–58.

Chomsky, N. (1995). Minimalist program. MIT Press, Cambridge, MA.

Cohen, M. X. (2017). Where does EEG come from and what does it mean? Trends in Neurosciences, 40(4):208–218.

Craver, C. F. (2007). Explaining the brain: mechanisms and the mosaic unity of neuroscience. Clarendon Press-Oxford University Press, New York, NY.

De Mulder, W., Bethard, S., and Moens, M.-F. (2015). A survey on the application of recurrent neural networks to statistical language modeling. Computer Speech & Language, 30(1):61–98.

Dehaene, S., Meyniel, F., Wacongne, C., Wang, L., and Pallier, C. (2015). The neural representation of sequences: from transition probabilities to algebraic patterns and linguistic trees. Neuron, 88(1):2–19.

Demberg, V. and Keller, F. (2008). Data from eye-tracking corpora as evidence for theories of syntactic processing complexity. Cognition, 109(2):193–210.

Devlin, J. T. and Watkins, K. E. (2007). Stimulating language: insights from TMS. Brain, 130(3):610–622.

Dubey, A., Keller, F., and Sturt, P. (2013). Probabilistic modeling of discourse-aware sentence processing. Topics in Cognitive Science, 5(3):425–451.

Embick, D. and Poeppel, D. (2015). Towards a computational(ist) neurobiology of language: correlational, integrated and explanatory neurolinguistics. Language, Cognition and Neuroscience, 30:357–366.

Fetsch, C. R. (2016). The importance of task design and behavioral control for understanding the neural basis of cognitive functions. Current Opinion in Neurobiology, 37:16–22.

Forstmann, B. U. and Wagenmakers, E.-J., editors (2015). An introduction to model-based cognitive neuroscience. Springer New York, New York, NY.

Forstmann, B. U., Wagenmakers, E.-J., Eichele, T., Brown, S., and Serences, J. T. (2011). Reciprocal relations between cognitive neuroscience and formal cognitive models: opposites attract? Trends in Cognitive Sciences, 15(6):272–279.

Frank, S. L. (2010). Uncertainty reduction as a measure of cognitive processing effort. In Proceedings of the 2010 Workshop on Cognitive Modeling and Computational Linguistics, pages 81–89, Uppsala, Sweden. Association for Computational Linguistics.

Frank, S. L. (2013). Uncertainty reduction as a measure of cognitive load in sentence comprehension. Topics in Cognitive Science, 5(3):475–494.

Frank, S. L. and Bod, R. (2011). Insensitivity of the human sentence-processing system to hierarchical structure. Psychological Science, 22(6):829–834.

Frank, S. L., Otten, L. J., Galli, G., and Vigliocco, G. (2015). The ERP response to the amount of information conveyed by words in sentences. Brain and Language, 140:1–25.

Frank, S. L. and Vigliocco, G. (2011). Sentence comprehension as mental simulation: an information-theoretic perspective. Information, 2(4):672–696.

Friederici, A. and Singer, W. (2015). Grounding language processing on basic neurophysiological principles. Trends in Cognitive Sciences, 19(6):1–10.

Friederici, A. D. (2012). The cortical language circuit: from auditory perception to sentence comprehension. Trends in Cognitive Sciences, 16(5):262–268.

Friederici, A. D. and Weissenborn, J. (2007). Mapping sentence form onto meaning: The syntax-semantic interface. Brain Research, 1146:50–58.

Gibson, E. and Pearlmutter, N. J. (1998). Constraints on sentence comprehension. Trends in Cognitive Sciences, 2(7):262–268.

Gibson, E. and Thomas, J. (1999). Memory limitations and structural forgetting: the perception of complex ungrammatical sentences as grammatical. Language and Cognitive Processes, 14(3):225–248.

Goodman, N. D. and Frank, M. C. (2016). Pragmatic language interpretation as probabilistic inference. Trends in Cognitive Sciences, 20(11):818–829.

Griffiths, T. L. (2011). Rethinking language: how probabilities shape the words we use. Proceedings of the National Academy of Sciences, 108(10):3825–3826.

Hagoort, P. (2009). Reflections on the Neurobiology of Syntax. In Bickerton, D. and Szathmáry, E., editors, Biological foundations and origin of syntax. MIT Press, Cambridge, MA.

Hagoort, P. (2013). MUC (Memory, Unification, Control) and beyond. Frontiers in Psychology, 4(July):416.

Hagoort, P. (2014). Nodes and networks in the neural architecture for language: Broca’s region and beyond. Current Opinion in Neurobiology, 28:136–141.

Hagoort, P. and van Berkum, J. (2007). Beyond the sentence given. Philosophical Transactions of the Royal Society, 362(1481):801–811.

Hale, J. (2001). A probabilistic earley parser as a psycholinguistic model. In Second meeting of the North American Chapter of the Association for Computational Linguistics on Language technologies 2001 - NAACL’01, pages 1–8, Morristown, NJ, USA. Association for Computational Linguistics.

Hale, J. (2003). The information conveyed by words in sentences. Journal of Psycholinguistic Research, 32(2):101–123.

Hale, J. (2006). Uncertainty about the rest of the sentence. Cognitive Science, 30(4):643–72.

Hale, J. (2011). What a rational parser would do. Cognitive Science, 35(3):399–443.

Hale, J. (2016). Information-theoretical complexity metrics. Language and Linguistics Compass, 10:397–412.

Hansen, P. C., Kringelbach, M. L., and Salmelin, R., editors (2010). MEG: An introduction to methods. Oxford University Press, New York, NY.

Hasson, U. and Honey, C. J. (2012). Future trends in neuroimaging: neural processes as expressed within real-life contexts. NeuroImage, 62(2):1272–1278.

Henderson, J. M., Choi, W., Lowder, M. W., and Ferreira, F. (2016). Language structure in the brain: A fixation-related fMRI study of syntactic surprisal in Reading. NeuroImage, 132:293–300.

Hickok, G. and Poeppel, D. (2007). The cortical organization of speech processing. Nature Reviews Neuroscience, 8(May):393–402.

Hochreiter, S. and Schmidhuber, J. (1997). Long short-term memory. Neural Computation, 9(8):1735–1780.

Huettig, F. (2015). Four central questions about prediction in language processing. Brain Research, (1626):118–135.

Huettig, F., Rommers, J., and Meyer, A. S. (2011). Using the visual world paradigm to study language processing: A review and critical evaluation. Acta Psychologica, 137(2):151–171.

Jackendoff, R. (2002). Foundations of language. Oxford University Press, New York.

Jonas, E. and Kording, K. P. (2017). Could a neuroscientist understand a microprocessor? PLOS Computational Biology, 13(1).

Jurafsky, D. (2002). Probabilistic modeling in psycholinguistics: linguistic comprehension and production. Probabilistic Linguistics, 30(1959):1–50.

Jurafsky, D. and Martin, J. H. (2009). Speech and language processing: an introduction to natural language processing, computational linguistics, and speech recognition. Pearson/Prentice Hall, Upper Saddle River, NJ, 2nd edition.

Kaan, E. and Swaab, T. Y. (2002). The brain circuity of syntactic comprehension. Trends in Cognitive Sciences, 6(8):350–356.

Krakauer, J. W., Ghazanfar, A. A., Gomez-Marin, A., MacIver, M. A., and Poeppel, D. (2017). Neuroscience needs behavior: correcting a reductionist bias. Neuron, 93(3):480–490.

Kuhl, P. K. (2010). Brain mechanisms in early language acquisition. Neuron, 67(5):713–727.

Kumaran, D., Hassabis, D., and McClelland, J. L. (2016). What learning systems do intelligent agents need? Complementary learning systems theory updated.

Kuperberg, G. R. (2007). Neural mechanisms of language comprehension: Challenges to syntax. Brain Research, 1146:23–49.

Kuperberg, G. R. and Jaeger, T. F. (2015). What do we mean by prediction in language comprehension? Language Cognition & Neuroscience, 3798(December 2015):1–70.

Kutas, M. and Federmeier, K. D. (2000). Electropsysiology reveals semantic memory use in language comprehension. Trends in Cognitive Science, 12(12):463–470.

Lau, J. H., Clark, A., and Lappin, S. (2017). Grammaticality, acceptability, and probability: a probabilistic view of linguistic knowledge. Cognitive Science, 41:1202–1241.

Lerner, Y., Honey, C. J., Silbert, L. J., and Hasson, U. (2011). Topographic mapping of a hierarchy of temporal receptive windows using a narrated story. The Journal of Neuroscience, 31(8):2906–2915.

Levinson, S. C. (2015). Turn-taking in Human Communication - Origins and Implications for Language Processing. Trends in Cognitive Sciences, 20(1):6–14.

Levy, R. (2008). Expectation-based syntactic comprehension. Cognition, 106(3):1126–1177.

Linzen, T. and Jaeger, F. (2014). Investigating the role of entropy in sentence processing. In Proceedings of the Fifth Workshop on Cognitive Modeling and Computational Linguistics, pages 10–18, Baltimore. Association for Computational Linguistics.

Logothetis, N. K. (2008). What we can do and what we cannot do with fMRI. Nature, 453(7197):869–78.

Lopopolo, A., Frank, S. L., van den Bosch, A., and Willems, R. M. (2017). Using stochastic language models (slm) to map lexical, syntactic, and phonological information processing in the brain. PLOS ONE, 12(5):1–18.

Luck, S. J. (2005). An Introduction to the Event-Related Potential Technique. MIT Press, Cambridge, MA.

Marcus, G., Marblestone, A., and Dean, T. (2014). The atoms of neural computation. Science, 346(6209):551–552.

Marcus, M., Marcinkiewicz, M., and Santorini, B. (1993). Building a large annotated corpus of English: the Penn Treebank. Computational Linguistics, 19(2):313–330.

Marr, D. (1982). Vision: a computational investigation into the human representation and processing of visual information. MIT Press, Cambridge, MA.

Mars, R. B., Shea, N. J., Kolling, N., and Rushworth, M. F. S. (2012). Model-based analyses: Promises, pitfalls, and example applications to the study of cognitive control. The Quarterly Journal of Experimental Psychology, 65(2):252–267.

Martin, A. E. (2016). Language processing as cue integration: grounding the psychology of language in perception and neurophysiology. Frontiers in Psychology, 7:120.

Nelson, M. J., Karoui, I. E., Giber, K., Yang, X., Cohen, L., Koopman, H., Cash, S. S., Naccache, L., Hale, J. T., Pallier, C., and Dehaene, S. (2017). Neurophysiological dynamics of phrase-structure building during sentence processing. Proceedings of the National Academy of Sciences, 114(18):E3669–E3678.

Norris, D., McQueen, J. M., and Cutler, A. (2000). Merging information in speech recognition: feedback is never necessary. Behavioral and Brain Sciences, 23(3):299–325; discussion 325–370.

Padó, U., Crocker, M. W., and Keller, F. (2009). A probabilistic model of semantic plausibility in sentence processing. Cognitive Science, 33(5):794–838.

Pallier, C., Devauchelle, A.-D., and Dehaene, S. (2011). Cortical representation of the constituent structure of sentences. Proceedings of the National Academy of Sciences, 108(6):2522–2527.

Palmeri, T. J., Love, B. C., and Turner, B. M. (2016). Model-based cognitive neuroscience. Journal of Mathematical Psychology, 76B(February):59–64.

Poeppel, D. (2012). The maps problem and the mapping problem: Two challenges for a cognitive neuroscience of speech and language. Cognitive Neuropsychology, 29(1–2):34–55.

Poldrack, R. A. (2010). Mapping mental function to brain structure: how can cognitive neuroimaging succeed? Perspectives on Psychological Science, 5(6):753–761.

Poldrack, R. A. and Gorgolewski, K. J. (2014). Making big data open: data sharing in neuroimaging. Nature Neuroscience, 17(11):1510–1517.

Pulvermüller, F., Garagnani, M., and Wennekers, T. (2014). Thinking in circuits: toward neurobiological explanation in cognitive neuroscience. Biological Cybernetics, 108(5):573–593.

Rayner, K. (1998). Eye movements in reading and information processing: 20 years of research. Psychological Bulletin, 124(3):372–422.

Roark, B. (2001). Probabilistic top-down parsing and language modeling. Computational Linguistics, 27(2):249–276.

Roark, B., Bachrach, A., Cardenas, C., and Pallier, C. (2009). Deriving lexical and syntactic expectation-based measures for psycholinguistic modeling via incremental top-down parsing. In Proceedings of the 2009 Conference on Empirical Methods in Natural Language Processing, volume 1, pages 324–333, Singapore. Association for Computational Linguistics.

Rosenfeld, R. (2000). Two decades of statistical language modeling: where do we go from here? Proceedings of the IEEE, 88:1270–1278.

Saffran, J. R. (2003). Statistical language learning. Current Directions in Psychological Science, 12(4):110–114.

Seidenberg, M. S. (1997). Language acquisition and use: Learning and applying probabilistic constraints. Science, 275(5306):1599–1603.

Small, S. L. and Nusbaum, H. C. (2004). On the neurobiological investigation of language understanding in context. Brain and Language, 89(2):300–311.

Smith, N. J. and Levy, R. (2011). Cloze but no cigar: The complex relationship between cloze, corpus, and subjective probabilities in language processing. Proceedings of the 33rd Annual Meeting of the Cognitive Science Conference, pages 1637–1642.

Staub, A. (2015). The effect of lexical predictability on eye movements in reading: critical review and theoretical interpretation. Language and Linguistics Compass, 9(8):311–327.

Stolk, A., Verhagen, L., and Toni, I. (2015). Conceptual alignment: how brains achieve mutual understanding. Trends in Cognitive Sciences, 20(3):180–191.

Taylor, W. L. (1953). ‘Cloze procedure’: a new tool for measuring readability. Journalism Quarterly, 30:415–433.

Van Schijndel, M. and Schuler, W. (2015). Evidence of syntactic working memory usage in MEG data. In Proceedings of the 6th Workshop on Cognitive Modeling and Computational Linguistics, pages 79–88, Denver, Colorado. Association for Computational Linguistics.

Wehbe, L., Ashish, V., Knight, K., and Mitchell, T. (2015). Aligning context-based statistical models of language with brain activity during reading. In Proceedings of the 2014 Conference on Empirical Methods in Natural Language Processing, pages 233–243, Doha. Association for Computational Linguistics.

Willems, R. M., editor (2015). Cognitive neuroscience of natural language use. Cambridge University Press, Cambridge.

Willems, R. M., Frank, S. L., Nijhof, A. D., Hagoort, P., and Van Den Bosch, A. (2016). Prediction during Natural Language Comprehension. Cerebral Cortex, 26(6):2506–2516.

Wu, S., Bachrach, A., Cardenas, C., and Schuler, W. (2010). Complexity Metrics in an Incremental Right-corner Parser. In Proceedings of the 48th Annual Meeting of the Association for Computational Linguistics, pages 1189–1198, Uppsala, Sweden. Association for Computational Linguistics.

Yarkoni, T., Speer, N. K., and Zacks, J. M. (2008). Neural substrates of narrative comprehension and memory. NeuroImage, 41(4):1408–1425.

